# A Novel Approach for Multi-Domain and Multi-Gene Family Identification Provides Insights into Evolutionary Dynamics of Disease Resistance Genes in Core Eudicot Plants

**DOI:** 10.1101/002766

**Authors:** Johannes A. Hofberger, Beifei Zhou, Haibao Tang, Jonathan D. G. Jones, M. Eric Schranz

## Abstract

Recent advances in DNA sequencing techniques resulted in more than forty sequenced plant genomes representing a diverse set of taxa of agricultural, energy, medicinal and ecological importance. However, gene family curation is often only inferred from DNA sequence homology and lacks insights into evolutionary processes contributing to gene family dynamics. In a comparative genomics framework, we integrated multiple lines of evidence provided by gene synteny, sequence homology and protein-based Hidden Markov Modelling to extract homologous super-clusters composed of multi-domain resistance (R)-proteins of the NB-LRR type (for NUCLEOTIDE BINDING/LEUCINE-RICH REPEATS), that are involved in plant innate immunity. To assess the diversity of R-proteins within and between species, we screened twelve eudicot plant genomes including six major crops and found a total of 2,363 *NB-LRR* genes. Our curated R-proteins set shows a 50% average for tandem duplicates and a 22% fraction of gene copies retained from ancient polyploidy events (ohnologs). We provide evidence for strong positive selection acting on all identified genes and show significant differences in molecular evolution rates (Ka/Ks-ratio) among tandem- (mean = 1.59), ohnolog (mean = 1.36) and singleton (mean = 1.22) R-gene duplicates. To foster the process of gene-edited plant breeding, we report species-specific presence/absence of all 140 *NB-LRR* genes present in the model plant *Arabidopsis* and describe four distinct clusters of *NB-LRR* “gatekeeper” loci sharing syntelogs across all analyzed genomes. In summary, we designed and implemented an easy-to-follow computational framework for super-gene family identification, and provide the most curated set of *NB-LRR* genes whose genetic versatility among twelve lineages can underpin crop improvement.

## INTRODUTION

To elucidate the dynamics underlying pathway and trait evolution across multiple lineages, it is of paramount importance to identify and distinguish the complete set of orthologous and paralogous loci present within multiple genome annotations in a phylogenetic framework (Lyons and Freeling 2008). Two homologous genes are referred to as orthologs if they descend from one locus present in the common ancestor lineage and diverged due to speciation (Fitch 1970). By definition, orthologous genes are embedded in chromosomal segments derived from the same ancestral genomic region, thus sharing high inter-species synteny between closely related lineages (Lyons et al. 2008). In contrast, paralogous loci refer to homologs within one lineage and are due to, for example, tandem-, transpositional- or whole genome duplications (WGDs) (Ohno 1970; Freeling 2009). Large-scale synteny is not observed for paralogs derived from small-scale events like tandem- and transpositional duplication. In contrast, paralogs derived from WGDs are located within intra-species syntenic genomic blocks, and can be referred to as ohnologs or syntelogs (Wolfe 2000; Bowers et al. 2003).

Increasing evolutionary distance between species presents limitations for homolog (ortholog and/or paralog) identification based on DNA sequence homology (such as blastn) due to the degeneration of the genetic code on the 3^rd^ codon position as well variable gaps of non-coding sequences (Wall et al. 2003; Alkatib et al. 2012). Hence, protein sequence identity and profile searches are used to infer homologs of protein-coding loci across distant clades with better sensitivity (Altschul et al. 1990; Krogh et al. 1994). Notably, proteins are organized in functional units termed domains (Vogel et al. 2004). For example, 37% of the *A. thaliana* Col-0 TAIR10 representative proteins contain more than one characterized protein domain (Swarbreck et al. 2008). Therefore, robust ortholog gene identification based on the encoded protein sequence often involves identification of multiple domain-specific sub-clusters followed by detection of overlaps to form a super-cluster composed of homologous multi-domain proteins.

Currently, there are several methods available for homolog detection. Among them, the determination of reciprocal best blast hits (RBH) at key phylogenetic nodes is the easiest and hence the most common method to infer families of orthologous and paralogous genes that represent the modern descendants of ancestral gene sets (used in, for example, (Wang et al. 2011a)). RBH comprise pairs of genes in two different genomes that are more similar to each other than either is to any other gene in the other genome. It is now evident that RBH-only approaches miss up to 60% of true orthologs in duplicate-rich species, which are particularly prevalent among angiosperm plants (Dalquen and Dessimoz 2013). Blasting Hidden Markov Model (HMM)-generated protein domain consensus sequences against a translated genome assembly is a more accurate way (used in, for example, (Guo et al. 2011)). The quality of results depends on the accuracy of the input sequences used to generate the consensus as well as the exonic structure of the proteins (i.e. some domains can span multiple small exons which can reduce the sensitivity of the translated blast). This creates challenges when analyzing highly-diverged gene families in more distant clades. In contrast, OrthoMCL (Li et al. 2003) employs a Markov Cluster algorithm to group putative orthologs and paralogs in all-vs.-all blast screen within and between given genome assemblies. Gene families defined by OrthoMCL therefore represent relatively compact and coherent clusters of similar proteins. However, OrthoMCL is unaware of the domain structure found in the family members and is computationally intensive and impracticable for application to dozens of genome annotations given a limited timeframe.

None of the similarity-based methods utilize information provided by the genomic context of the putative ortholog / paralog gene pairs. This is relevant because two members of the same gene family can evolve differently in two lineages to a degree that the ortholog in lineage A produces a RBH to the paralogs in lineage B and vice versa (Hofberger et al. 2013). This produces false ortholog assignments but can be clarified by scoring and weighting gene synteny evidence (Tang and Lyons 2012). In this study, we design and employ a novel, iterative approach by combining blast, HMM modeling and genomic contextual information provided by synteny to identify robust multi-domain super-clusters. We do so by analyzing a total of 12 core eudicot genome annotations and identifying all present R-proteins of the NB-LRR type that convey critical functions in plant innate immunity against pathogens. In this context, we acknowledge an error-rate due to incomplete gene annotations, as shown for tomato and potato (Jupe et al. 2013) (Andolfo et al. 2014, submitted for publication). However, this error-rate (a) affects all aforementioned methods for gene clustering, (b) will decrease with the continued onset of whole genome (re)sequencing in the future and (c) can be minimized by application of scaffolds instead of gene models to our pipeline (see discussion).Plants have evolved a two-layered innate immune system against microbial and other pathogens (Jones and Dangl 2006). In a first layer of defense, transmembrane pattern recognition receptors (PRRs), usually with extracellular LRR-type domains, recognize pathogen associated molecular patterns (PAMPs) and initiate downstream signaling events including defense gene induction (Chinchilla et al. 2006), and lead also to cell wall reinforcement by callose deposition and SNARE-mediated secretion of anti-microbial compounds (Collins et al. 2003; Zipfel and Robatzek 2010). This is referred to as PAMP- or pattern-triggered immunity (PTI).

Successful pathogens have evolved apoplastic or cytoplasmic virulence factors (effectors) that overcome PTI (Schwessinger and Zipfel 2008). As a second layer of the innate immune response, many host plant lineages evolved intracellular R-proteins of the NB-LRR type that respond to virulence factors, either directly or through their effects on host targets (Hann et al. 2013). Plants producing a specific R-gene product are resistant towards a pathogen that produces the corresponding effector gene product (avirulence factors encoded by *Avr* genes), leading to gene-for-gene resistance (Van der Biezen and Jones 1998). This is referred to as effector-triggered immunity (ETI). Rounds of ETI and effector-triggered susceptibility (ETS) due to novel *Avr* genes on the pathogen side can result in an evolutionary arms-race, generating a “zigzagzig” amplitude of host resistance and susceptibility (Jones and Dangl 2006). R-genes play a major role in defending crops against microbial infection and thus are of great interest in plant breeding program and efforts to meet increased global food production. In potato, for example, R-proteins of the NB-LRR type confer resistance to the oomycete *Phytophthora infestans*, a hemibiotrophic oomycete pathogen that causes late blight (Ballvora et al. 2002; Vleeshouwers et al. 2011). In *Arabidopsis*, R-proteins of the NB-LRR type have been studied extensively in terms of molecular function, structural organization, sequence evolution and chromosomal distribution (Botella et al. 1998; Young 2000; Meyers et al. 2003; Guo et al. 2011). This superfamily is encoded by scores of diverse genes per genome and subdivides into TIR-domain-containing (for TOLL / INTERLEUKIN LIKE RECEPTOR/RESISTANCE PROTEIN) (TIR-NB-LRR or TNL) and non-TIR-domain containing (NB-LRR or NL), including coiled-coil domain-containing (CC-NB-LRR or CNL) R-protein subfamilies (McHale et al. 2006). For example, the TNL type R-protein RPP1 confers resistance to *Hyaloperonospora arabidopsidis* (downy mildew) in *Arabidopsis* (Boisson et al. 2003). Similarly, the RPS5 CNL type R-protein interacts in a gene-for-gene relationship with the avrPphB effector from *Pseudomonas syringae* to activate innate immune responses (Warren et al. 1998). The TNL type R-protein RRS1 confers resistance to the soil microbe *Ralstonia solanacearum* in *Arabidopsis* (Deslandes et al. 2003). The latter also contains a C-terminal WRKY transcription factor-like domain for DNA binding (Bernoux et al. 2008), increasing the number of domains common to NB-LRR clusters to five. This number is further extended by cases with presence of variable N-terminal domains mediating extended gene function. For example, the *Arabidopsis NB-LRR* type locus *CHILLING-SENSITIVE3* (*CHS3*) encodes a mutated allele of an N-terminal LIM-type domain-carrying TNL protein, leading to constitutive activation of defense responses and decreased chilling susceptibility in *35S::CHS3* overexpression lines (Yang et al. 2010).

TIR- and non-TIR NB-LRR protein clusters share a conserved central NB-ARC domain including three subdomains (NB, ARC1, and ARC2). Together, these confer ATPase function (van Ooijen et al. 2008). The C-terminal part of NB-LRR proteins harbors a leucine-rich repeat (LRR)-domain for recognition of intracellular effector molecules upon infection, leading to a conformational shift within the NB-ARC domain (Boller and Felix 2009) upon recognition of the corresponding effector or a change in the surveyed plant protein. In case of the soybean (*Glycine max*) CNL-class R-protein RPSk-1, DNA binding affinity changes upon *Phytophthora sojae* effector recognition. As a consequence the R-protein can act as transcription factor after diffusion to the nucleus to induce defense gene induction (Tameling et al. 2006; Bhattacharyya 2007; Takken and Goverse 2012).

A genome-wide comparison of multi-gene families in *A. thaliana* Col-0 revealed a strong contribution of gene duplication to the *NB-LRR* gene count and genomic distribution (Cannon et al. 2004). For example, 63% of all reported *NB-LRR* genes are members of tandem arrays in both *A. thaliana* (101 / 159) and *A. lyrata* (118 / 185) (Guo et al. 2011). Notably, *NB-LRR* loci are subject to positive selection (Mondragón-Palomino et al. 2002). In this context, (Guo et al. 2011) re-assessed rates of molecular evolution for both sets of tandem- and non-tandem (singleton hereafter) and found significant differences in selection rates. In this study, we went a step further by distinguishing the contribution of tandem- and syntelog-duplicates to *NB-LRR* cluster expansion and diversity within a wider phylogenomics perspective, thereby covering an evolutionary timeframe of approximately 100 MA that corresponds to the radiation of core eudicots (Bremer et al. 2009; Jiao et al. 2012). We compared the average rates of molecular evolution for singleton-, tandem- and syntelog duplicate R-genes. We further provide evidence for strong positive, but significantly different, selection rates acting on all copy classes of *NB-LRR* duplicates, illustrating the impact of gene and genome duplication to the diversification of plant key traits across approximately 100 MA of genome evolution.Recent genomic analyses have revealed a common history of ancient, successive polyploidy events that are a common feature of genome evolution shared by all flowering plant linages (Lyons et al. 2008). For example, the *Arabidopsis* lineage underwent at least five polyploidy events that we know of, two preceding and three following angiosperm radiation (Jiao et al. 2011). The most recent WGD event for the *Arabidopsis* lineage is termed At-α and shared by all Brassicaceae including the extant sister clade Aethionemeae (Schranz et al. 2012; Haudry et al. 2013). The older At-β WGD is shared by most species in the order Brassicales, but occurred after the papaya lineage split (Ming et al. 2008; Barker et al. 2009). The more ancient At-y event is a whole genome triplication (WGT) that is shared by most eudicots including all Rosids, all Asterids (including tomato), Grape (Vitales) and more distant and basal clades such as *Gunnera manicata* (Gunnerales) and *Pachysandra terminalis* (Buxales) (Jaillon et al. 2007; Vekemans et al. 2012). In addition to ancient polyploidy events, more recent, species-specific WGDs/WGTs occurred in various lineages, such as genome triplications in *B. rapa* (Tang and Lyons 2012) (Br-α WGT), *T. hasslerania* (Th-α WGT) (Barker et al. 2009; Cheng et al. 2013) and the Solanaceae (Tomato Genome Consortium 2012). Hence, the “syntenic depth” (defined as the level of genome multiplicity expected from the multiplication of successive WGDs/WGTs) of the *Brassica rapa* genome is 36x compared to the putative 1x eudicot ancestor (3x due to At-y, 2x more due to At-β, 2x more due to At-α and finally 3x due to Br-α). Under consideration of two polyploidy rounds at or near the origin of angiosperms as well as 2x at or near the origin of seed plants (Jiao et al. 2011), the syntenic depth of the *B. rapa* genome would be expected to be increased to 144x (“rho-mu-delta-ploidy” genome).

Both recent and ancient polyploidy results in pairwise syntenic regions scattered throughout the replicated genome, defined as consecutive copies of ohnologs and remnant patterns of (ancient) polyploidy (genomic blocks) (Bowers et al. 2003). Genome duplication is often followed by a genome-wide process of biased fractionation that preferentially targets one sub-genome to retain clusters of dose-sensitive genes often organized in functional modules (Freeling and Thomas 2006; Thomas et al. 2006; Schnable et al. 2011). Recently, evidence is accumulating for the connection of ancient WGD events to birth and diversification of key biological traits. In Brassicaceae, WGD shaped the genetic versatility of the glucosinolate pathway (Hofberger et al. 2013), a key trait mediating herbivore resistance and thus highly connected to reproductive fitness of the population. Similarly, starch biosynthesis in grasses, origin and diversification of seed and flowering plants as well as increased species survival rates on the KT-boundary are hypothesized to be linked to ancient polyploidy events (De Bodt et al. 2005; Irish and Litt 2005; Fawcett et al. 2009; Paterson et al. 2010; Jiao et al. 2012).

Polyploidy events also influence other kinds of duplication, thereby creating a network of factors with mutual influence. In *Brassica rapa* (that underwent an additional species-specific genome triplication event, see above), arrays of tandem duplicate (TD) genes (TAR genes) fractionated dramatically after the Br-α WGT event when compared either to non-tandem genes in the B. rapa or to tandem arrays in closely related species that have not experienced a recent polyploidy event (Fang et al. 2012). Errors in DNA replication due to template slippage or unequal crossing-over can result in tandem duplication (TD), producing tandem arrays (TAR) of paralog genes in close genomic proximity (Kane et al. 2010). It is known that TAR genes are enriched for genes functioning in biotic and abiotic stress (Rizzon et al. 2006). For disease resistance, there are multiple taxa with an evident impact of TD to trait evolution, including members of Brassicaceae (Leister 2004), Solanaceae (Parniske et al. 1999) and Fabaceae (Bellieny-Rabelo et al. 2013). In this study, we determine the fraction of tandem- and whole genome duplicate copies among all (re)annotated full-length *NB-LRR* genes across 12 species in the context of a phylogenomics perspective, based on uniform standards facilitating all comparisons. After utilization of duplicate classes, we assess and compare rates of molecular evolution to describe a complex interplay of TD and WGD events driving R-protein super-family extension, both of which expanded the evolutionary playground for functional diversification and thus potential novelty and success.

In contrast to near-exponential growth of the human population, access to arable land is limited and often diminishes irretrievable resources (such as rainforest) when increased (Walsh et al. 2011). Genetic modification (GM) of crop plants to increase yields with the remaining land has therefore become a well-established strategy for feeding the planet without destroying it. For example, global GM crop plantings increased 100-fold from 1996 to 2012 (Clive 2012). However, GM crops have been the subject of intense debate due to concerns about hypothetical environmental and health implications of transgenics (Gaskell et al. 1999; Waigmann et al. 2012). In response to consumers’ reticence, new techniques are now being developed that are faster than traditional breeding methods but notably do not involve the introduction of foreign DNA. Such techniques commonly employ targeted mutagenesis and are referred to as (plant) genome editing or targeted (plant) genome engineering (Belhaj et al. 2013). To date, three classes of bacterial nucleases can be engineered by customized small RNAs to introduce DNA double-strand breaks (DSBs) at targeted sites within the plant genome, followed by subsequent elicitation of the plant's endogenous toolbox to repair the induced breaks by natural processes of homologous recombination (HR) or non-homologous end-joining (NHEJ). Zinc finger nucleases (ZFNs), TAL effector nucleases (TALENs) as well as Cas9 endonuclease all can generate single or multiple gene knock-out lines as well as plants carrying single-nucleotide polymorphisms (SNPs) in targeted genes) (Boch 2011; Carroll 2011; Hsu et al. 2013). Notably, none of the resulting plants can be identified as GM due to the absence of marker genes or other foreign DNA fragments (Belhaj et al. 2013). In the context of *NB-LRR* type R-gene products, single SNPs in combination with otherwise low nucleotide diversity are evident to confer resistance to extended groups of pathogens (Bakker et al. 2006; Jermstad et al. 2006; Liu and Ekramoddoullah 2007; Hayashi et al. 2010). In this study, we provide an overview of site-specific nucleotide diversity among the complete set of full-length *NB-LRR* genes present in 12 eudicot genome annotations (including 6 major crops). Likewise, we provide a matrix referencing both data on *NB-LRR* gene sequence and function evident in *A. thaliana* to any of the other analyzed genomes. We thereby seek to facilitate design of future experiments for generation of gene-edited crops, ultimately necessary to meet global food production in an ever-increasing population unwilling to rely on GM plants.

## RESULTS

### Determination of protein domain-specific clusters

Encoded architecture of *NB-LRR* loci in plants is variable and can comprise up to 5 different domains in *Arabidopsis* (**Fig. 1**). In contrast to previous studies (Meyers et al. 2003), we define functional *NB-LRR* proteins as composite units sharing both NB-ARC domain and a LRR domain signal due to at least one repeat. Hence, TIR-NB-, LRR-only, NB-only or TIR-only proteins are not assigned as *NB-LRR* proteins by definition. To determine the number of *NB-LRR* loci within a given genome annotation, we combine layers of information provided by sequence homology, protein identity as well as genomic context of target genes in a custom, iterative approach using batch programming (**Fig. 2**).

**Figure 1.**
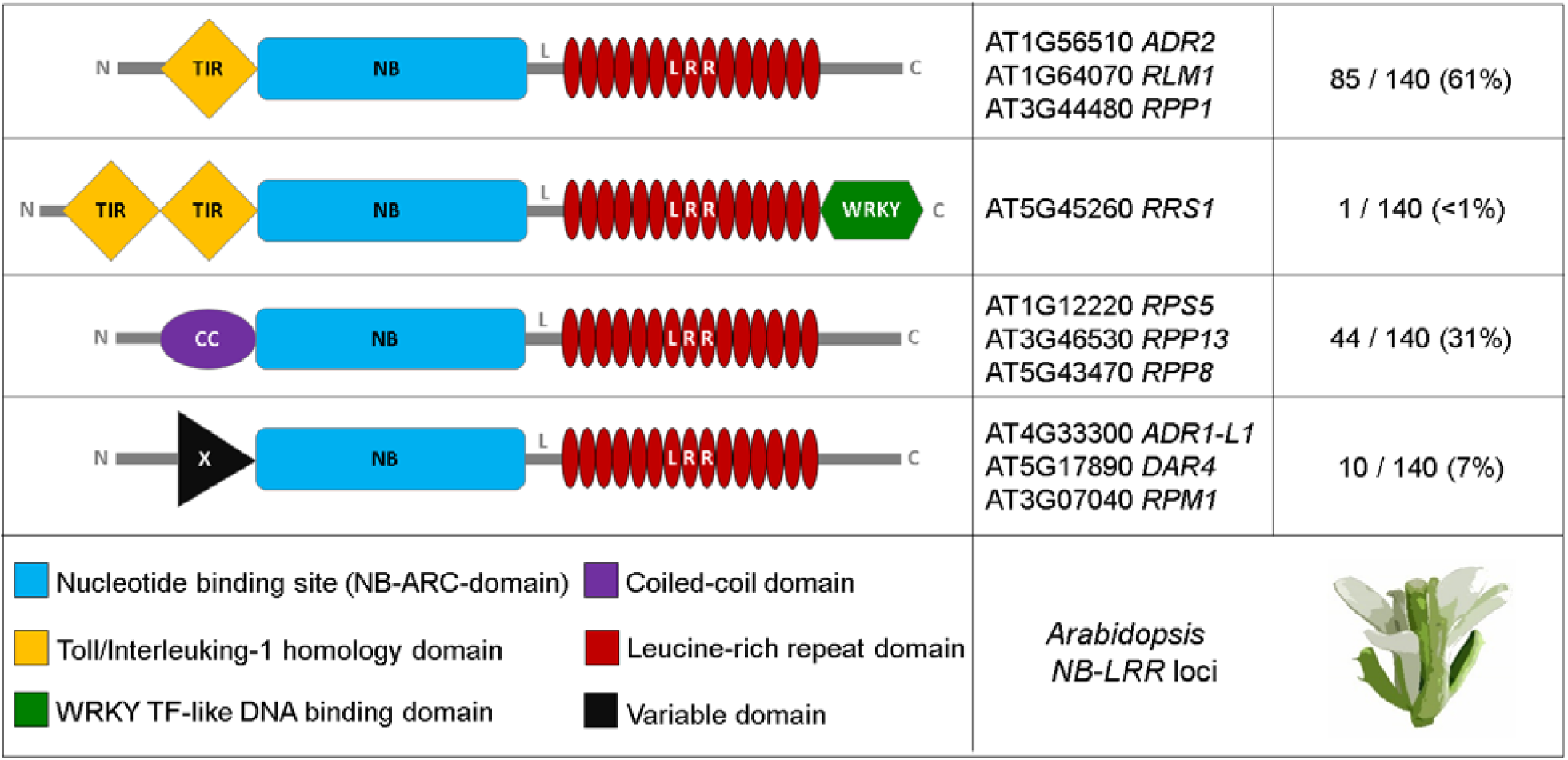
The *NB-LRR* multi-gene family comprises more than five domain classes. Four gene clusters in *Arabidopsis* are defined in this study based on domain compositions. Left: Frequent domain combinations. Middle: Well-characterized class representative in *Arabidopsis thaliana* Col-0. Right: relative abundance of target *NB-LRR* locus class in *Arabidopsis thaliana* Col-0.

**Figure 2.**
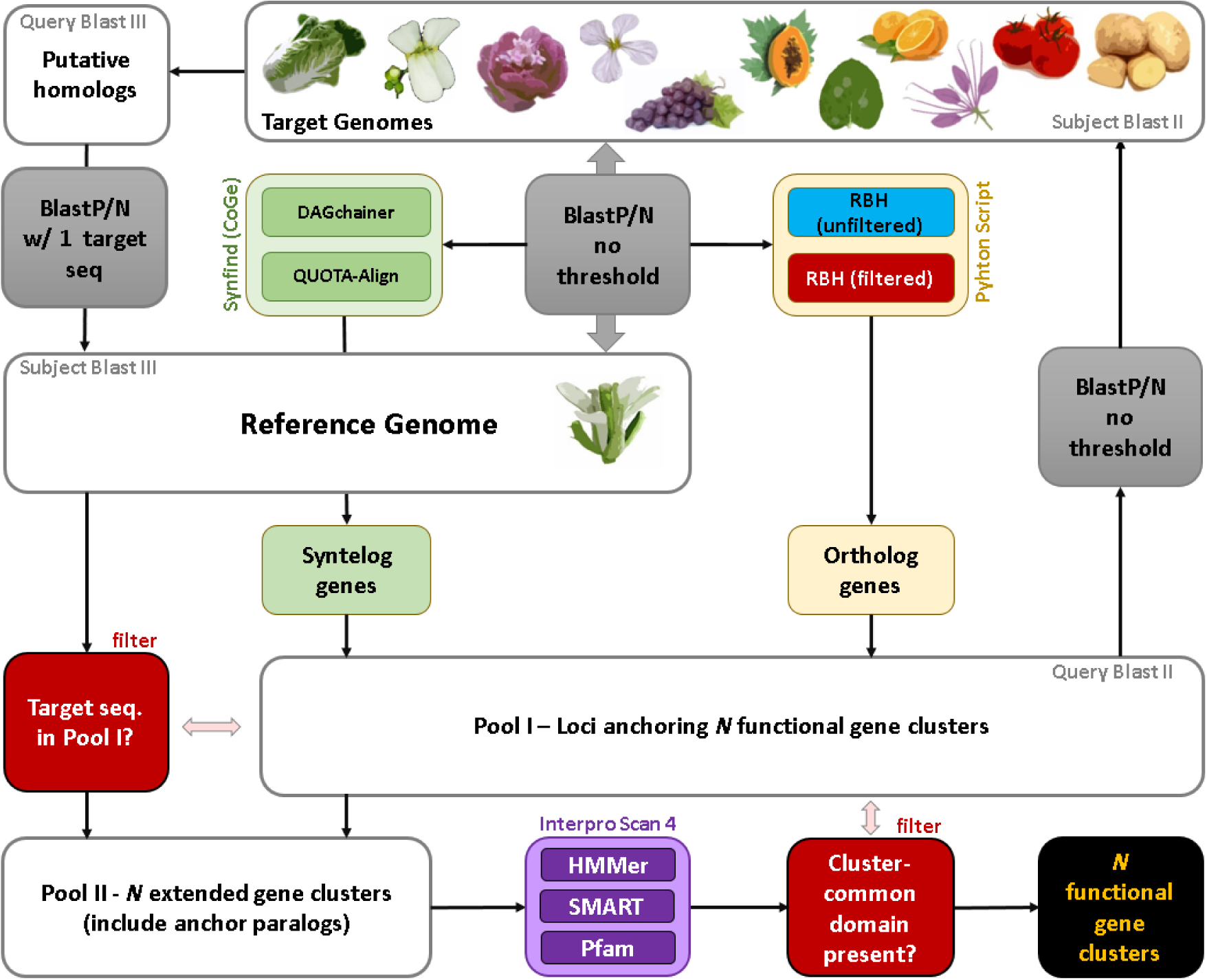
Custom flowchart showing the integrated approach for the identification of conserved multi-domain protein clusters. A bi-directional blast screen between a reference genome and n target genomes marks the entry point to the pipeline (grey box in the upper middle). Grey indicates blast screens. Red indicates filtering steps. White boxes indicate pools of FASTA-formatted sequences. Purple indicates Hidden Markov Modelling steps to predict and map protein domains. Green refers to the CoGe system (see methods). Ochre indicates custom python scripts. Flowchart starts with two-per-genome bidirectional blast screens (middle) and ends with highly accurate functional protein clusters (black, bottom right).

In the first step, we identified putative orthologous (based on RBH) and/or syntenic (based on Synfind) “anchor” genes (a) present in the most up-to-date genome annotations of (1) *A. lyrata*, (2) *B. rapa*, (3) *E. parvulum*, (4) *Ae. arabicum*, (5) *T. hasslerania*, (6) *C. papaya*, (7) *C. sinensis*, (8) *V. vinifera*, (9) *N. benthamiana*, (10), *S. lycopersicum* and (11) *S. tuberosum* as well as (b) aligning to any gene present in the (12) *A. thaliana* Col-0 TAIR10 genome annotation. This step resulted in a cluster dataset anchoring every gene family present in *Arabidopsis* to all of the aforementioned lineages, hence providing valuable means for gene identification with any kind of target trait known in core-eudicot plants (**supplementary table 1**). Subsequently, we screened for genes encoding (i) a LRR-domain, (ii) a NB-domain or (iii) a TIR-domain (extended set of target genes defined in this study, see methods) (**Fig. 2**, **supplementary table 1**). In a second step, we screened for anchor gene paralogs present in every aforementioned genome annotation to form an extended cluster of homologous genes containing at least one of the aforementioned domains (**Fig. 2**). In a third step, we applied multiple machine learning methods (see methods) to filter false-positives to obtain three highly accurate, functional domain cluster (NB-ARC/LRR/TIR) (**Fig. 2**, **supplementary table 2**). We performed the third (filtering) step three times (once for every aforementioned domain). For genome annotation versions and species-wise domain cluster gene count, see **fig. 3**.

**Figure 3.**
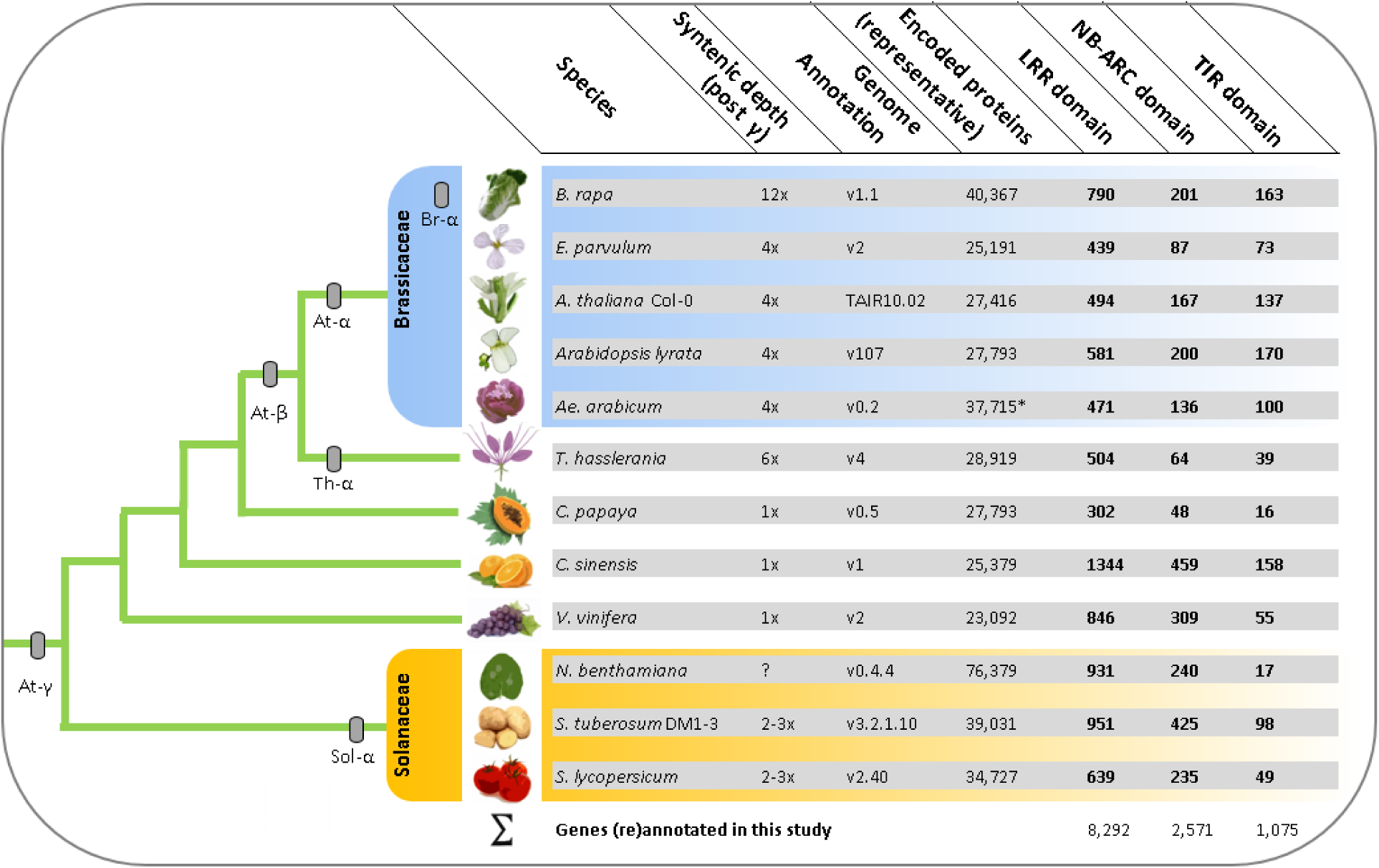
Results overview of plant Resistance (R)-gene domain (re)annotation. Shown left are phylogenetic relationships among all analyzed plant lineages and rough placement of whole-genome duplication events. Shown right are numbers of taget genes per domain cluster and information on annotaion build.

We identified 8,292 loci coding for a LRR-domain. Among those, the lowest number of genes containing a LRR domain is 302 for the *C. papaya* genome annotation v0.5. In contrast, the highest number of genes encoding a LRR domain is 1,344 for the *C. sinensis* genome annotation v1. Interestingly, both annotations share a syntenic depth of 2n, representing the lowest-copy genomes subjected to our analysis (i.e. no major evidence for WGD since At-γ).

We found 2,571 genes in total containing a NB-ARC domain across the twelve genomes. Likewise, the lowest number was found within the *C. papaya* genome annotation v0.5 (48 loci). Again, the highest number of genes encoding a NB-ARC domain was found in the *C. sinensis* genome annotation v1 (459 loci).

For genes encoding at least one TIR-domain, we identified a pool of 1,075 genes across the twelve genomes. Similar to the aforementioned cases, the *C. papaya* genome annotation v0.5 encodes the lowest number of TIR-like loci (16 genes). In contrast to the aforementioned cases, the *A. lyrata* annotation v1.07 (but not *C. sinensis*) contains the highest number of encoded TIR-domains (170 loci). Notably, the syntenic depth of *A. lyrata* is double that of papaya or orange. For phylogenetic relationships and syntenic depth levels of all analyzed genome annotation, see **fig. 3**.

### Determination of *NB-LRR* multi-gene family size

For every analyzed plant species, we determined the multi-gene family size of all annotated *NB-LRR* candidate genes by overlaying each filtered domain-clusters (**Fig. 4**). Note that statements about target loci missing or flawed within the gene annotations are beyond the scope of this section, but can likewise be considered *in silico* by applying sequence scaffolds/contigs instead of gene models to our customized pipeline (see discussion).

**Figure 4.**
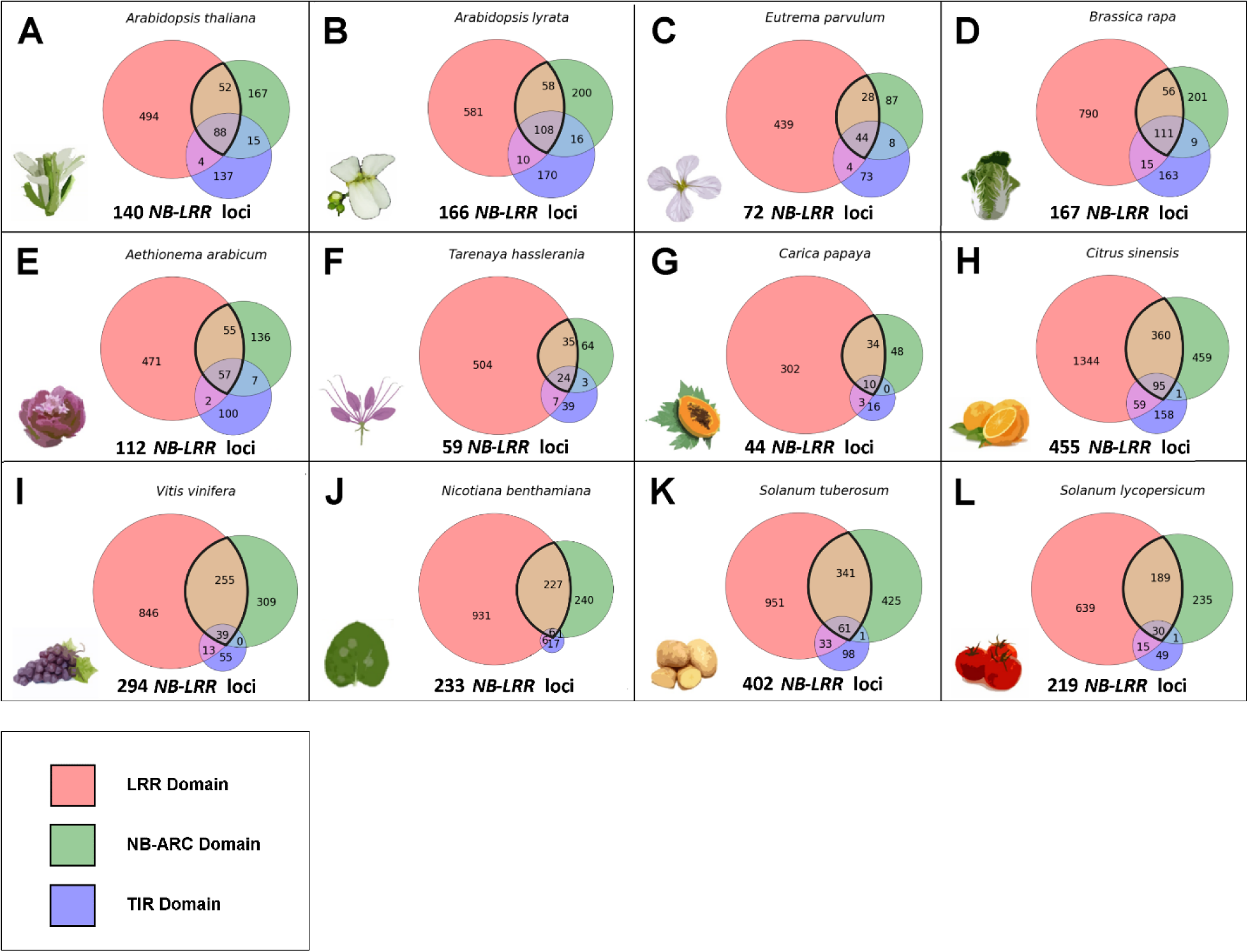
Area-weighted Venn diagrams illustrating the distribution of three main functional domains common to *NB-LRR* gene clusters shown for twelve species. Please note that we require an *NB-LRR* gene to harbor both NB-ARC and LRR domains.

For the *A. thaliana* Col-0 TAIR10 genome annotation, we have found 140 non-redundant *NB-LRR* loci (**Fig. 4A**). Previous studies found 166 (Richly et al. 2002), 178 (Meyers et al. 2003), 174 (Yang et al. 2008; Chen et al. 2010), and 138 (Guo et al. 2011) *NB-LRR* loci, respectively. In contrast, TAIR10 domain annotation efforts reported 127 target loci (Swarbreck et al. 2008). The differences in our study resulted from usage of the updated TAIR10.02 annotation and more stringent criteria; namely the exclusive combination of machine learning with sequence identity and consideration of the genomic context (e.g. synteny). For example, we focus on protein-coding genes only and ignore non-functional (i.e. pseudogenized) loci due to the scope of this study to provide information relevant for breeding of gene-edited crops. Moreover, we define *NB-LRR* proteins as sharing both NB and LRR domains, whereas many previous studies score anything as a *NB-LRR* gene that partially aligns to any one domain common to the cluster (i.e. TIR-only, NB-only, LRR-only genes).

For the *A. lyrata* genome annotation v0.2, we identified 166 non-redundant *NB-LRR* loci (**Fig. 4B**). Previous studies reported 182 (Chen et al. 2010) and 138 (Guo et al. 2011) *NB-LRR* loci, respectively. Chen et al. score pseudogenes as well as loci that don't contain both NB and LRR domain, leading to the higher number of target genes than reported in this study. The difference between our results and those of Guo et al. is likely due to false-negative target genes with a divergence level that cannot be recognized by their applied HMM-generated NB-consensus sequence. We were able to score these more divergent loci using synteny data to anchor locus determination and subsequent *de novo* domain prediction using a combination of 14 HMM algorithms (see methods) (**Fig. 2**). For example, the *A. lyrata* locus fgenesh1_pg.C_scaffold_8000651 displays only moderate homology (e-value: 1e-34) to its closest related sequence in *A. thaliana*, a P-loop containing nucleoside triphosphatase that is not defined as *NB-LRR* locus. However, we found both NB-ARC and LRR domain within that gene in *A. lyrata*.

For the crop plant *B. rapa* (genome annotation v1.1), we found 167 non-redundant *NB-LRR* candidate genes (**Fig. 4D**), while previous studies reported a sum of 92 (Mun et al. 2009). (Mun et al. 2009) did not have the whole genome assembly available, and hence identified R-proteins based on 1,199 partially redundant BAC clones mostly from a single chromosome. The authors acknowledge a significant degree of sequence redundancy within the available dataset, that covers 19-28% of the *B. rapa* genome only. Likewise, (Mun et al. 2009) perform *ab-initio* gene annotation based on the fgenesh algorithm only (Salamov and Solovyev 2000), and solely use protein sequence homology (based on blastp) for R-protein homolog identification. In contrast, we use the whole gene-space assembly (including every to-date annotated protein-coding gene) as well as three layers of information for homolog identification (see methods).

To our knowledge, we provide the first analyses of R-proteins for *E. parvulum*, *Ae. arabicum*, *T. hasslerania* and *N. benthamiana*. For the extremophile saltwater cress *E. parvulum* (previously known as *Thelungiella parvula*, genome annotation v2), we found 72 non-redundant *NB-LRR* loci (**Fig. 4C**). For *Ae. arabicum*, the extant sister lineage to all other mustard family members (genome annotation v0.2), we identified 112 non-redundant *NB-LRR* loci (**Fig. 4E**). For the *T. hasslerania* genome annotation v4 (previously known as *Cleome spinosa*), we identified 59 non-redundant *NB-LRR* loci for this species (**Fig. 4F**), that underwent a lineage-specific genome triplication (**Fig. 3**) and has been established as the mustard family outgroup (Schranz and Mitchell-Olds 2006; Barker et al. 2009; Cheng et al. 2013). For the Solanaceae and tobacco relative *N. benthamiana*, we identified 233 non-redundant *NB-LRR* proteins (**Fig. 4J**). Notably, *N. benthamiana* is widely used as system for transient over-expression and silencing of various genes involved in plant innate immunity to elucidate downstream signaling events after PAMP-mediated priming. In this context, our results provide mapping of all characterized *NB-LRR* mediated functions present in *A. thaliana* to the *Nicotiana* gene-space assembly (**supplementary table 2**), thereby facilitating adjusted planning of aforementioned experiments and better understanding of the results in the Solanaceae.

For the crop plant *C. papaya* (genome annotation v0.5), we identified 44 non-redundant R-proteins of the *NB-LRR* type (**Fig. 4G**). Among all species we have analyzed so far, the papaya gene-space assembly encodes the lowest number of R-gene candidates. We again acknowledge the possibility of incomplete gene annotations in this context (see discussion). However, the low gene count of the *NB-LRR* locus family was previously revealed for the available papaya genes set (Porter et al. 2009). The authors found 54 target loci using a combination of tblastx (Altschul et al. 1990) as well as the pfam HMM algorithm to search for the pfam NB (NB-ARC) family PF00931 domain (Bateman et al. 2004). The difference in gene-family size estimates is due to an updated genome annotation we have used, as well as more stringent criteria for target gene scoring (i.e. *NB-LRR* proteins are defined as sharing both NB- and LRR domain, see above).

Our analysis revealed 455 non-redundant loci of the *NB-LRR* type for the crop plant *C. sinensis* (orange) (**Fig. 4H**). Evidence for the high R-gene count in orange has been noted previously. For example, the plant resistance gene database (prgdb) lists 3,230 R-genes (including LRR-domain containing receptor-like kinases/proteins) for this crop plant (Hermoso et al. 2009), many of which are redundant. To our knowledge, our study comprises the first efforts to cross-reference both NB-ARC and LRR domains among R-genes in orange.

For grape (*V. vinifera*), we found 294 non-redundant R-proteins sharing both NB-ARC and LRR domains (**Fig. 4I**). Previous efforts identified 300 target genes (Yang et al. 2008). The differences are due to an updated genome assembly as well as more stringent criteria for *NB-LRR* locus definition.

In addition, we subjected the potato crop (*S. tuberosum* Group Phureja DM1-3) genome annotation v3.2.10 to our customized pipeline for identification of homologous gene clusters. We identified 402 encoded non-redundant *NB-LRR* proteins within the potato genome (**Fig. 4** K). Previous efforts identified 438 target genes (Jupe et al. 2012) from the annotated proteins using the MEME and MAST algorithms (Bailey et al. 2009) as well as 755 target genes based on reduced representation analysis of DNA enriched for the *NB-LRR* gene repertoire (Jupe et al. 2013). Referring to the latter study, we acknowledge the inability of our pipeline to identify genes present in the crop but flawed or missing from the annotation or the assembly. The difference between our value and (Jupe et al. 2012) results from more stringent criteria in *NB-LRR* locus identification. For example, at least 34 of the 438 genes from (Jupe et al. 2012) do not contain both NB-ARC and LRR domains, whereas at least two don't contain any of the required domains.

For tomato (*Solanum lycopersicum* Heinz 1706), we have found 219 non-redundant R-proteins of the *NB-LRR* type (**Fig. 4L**). Previous studies identified 221 target genes sharing both NB and LRR domains in a very conclusive approach (Andolfo et al. 2013). The minor difference in numbers is due to a different build of the annotation based on the genome version 2.4 (fusion of loci/locus fragments) and illustrates the thoroughness of the corresponding authors work. In total, we identified 2,363 R-proteins of the NB-LRR type. All CDS sequences are appended including translation to protein sequences. (supplementary data 3 and 4).

### *NB-LRR* gene localization and determination of tandem duplicate fractions

We localized all reported *NB-LRR* loci onto the corresponding chromosomes/scaffolds/contigs present in all analyzed genome assemblies except *N. benthamiana* (excluded from Circos plot due to insufficient assembly quality, see methods). Application of a number of n = 10 allowed gene spacers (see methods) allowed determination of a global rate of 53% tandem duplicates (**Fig. 5**). Notably, we have found significant differences in tandem array fractions between the analyzed species (up to a factor of 2.8). For example, 70% among all *NB-LRR* genes present in the *V. vinifera* genome annotation v2 are members of tandem arrays. In contrast, the *N. benthamiana* genome annotation v0.4.4 contains 25% tandem duplicates among all present *NB-LRR* loci. The latter represents a fragmented gene-space rather than a genome assembly, leading to a likely under-estimation of tandem duplicates fraction. Hence, the global tandem duplicates fraction drops to 50% after inclusion of *N. benthamiana* loci. For the mean gene count per *NB-LRR* tandem arrays, Aethionema scores highest with an average of 3.4. Likewise, the extant mustard family sister clade contains the largest tandem array we found so far, with a total of 11 genes. In contrast, the largest array in orange (*C. sinensis*) contains 5 genes only, leading to a genome-wide average of 2.2 *NB-LRR* genes per tandem array (Table 1). Please note that we require presence of both NB-ARC and LRR domains fur *NB-LRR*-type R-gene curation. Therefore, some of the aforementioned tandem arrays may be further extended due to the presence of partial sequences in close proximity. We do not exclude a biological significance of such fragments per se, but set the scope to full-length candidate genes exclusively for obtaining a uniform dataset to facilitate comparisons of molecular evolution rates (see below).

**Figure 5.**
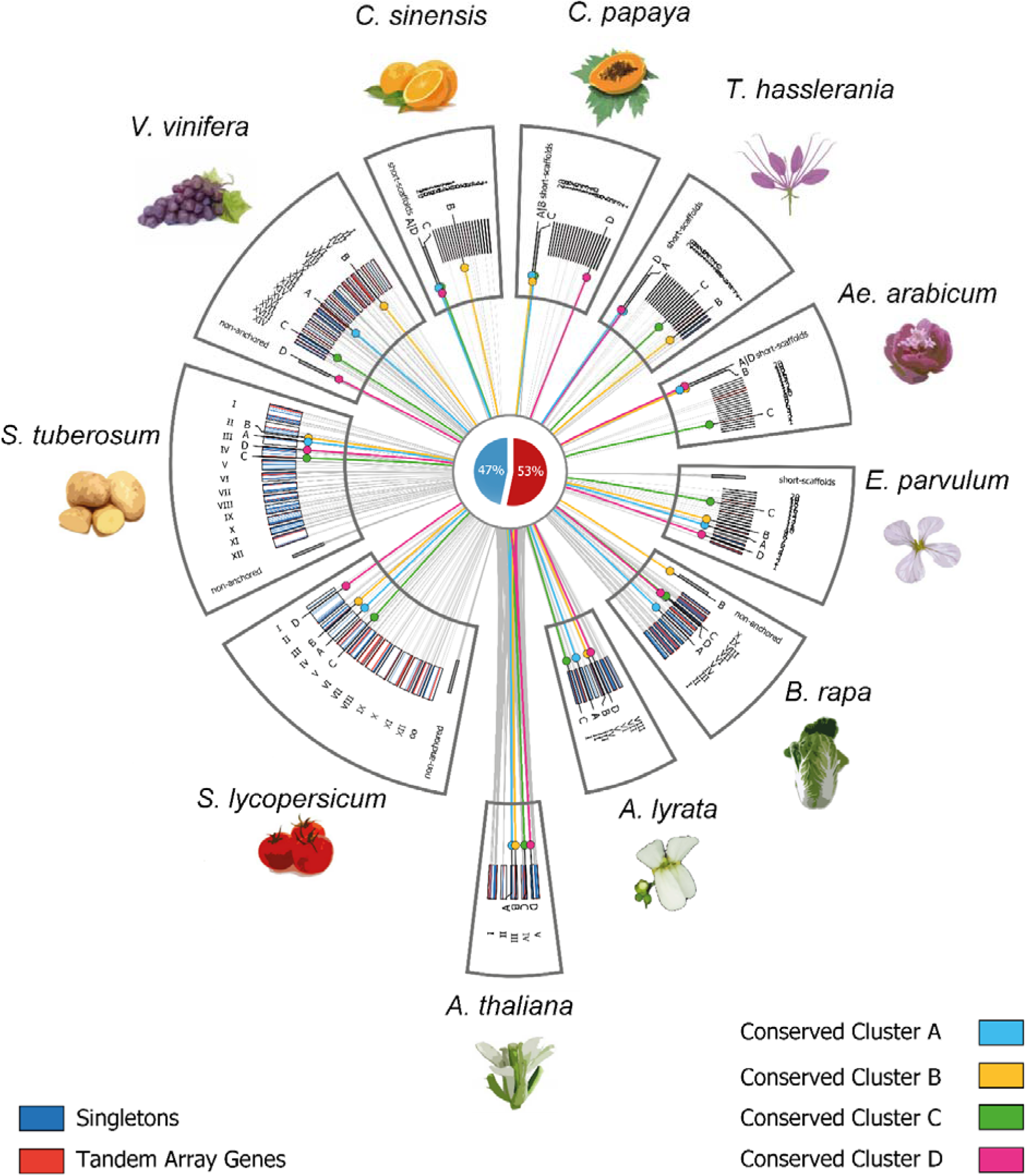
Circos ideogram with 2,363 *NB-LRR* loci localized on 11 genome annotations. Beginning at the bottom block in counter-clockwise orientation, shown are (1) *Arabidopsis thaliana* Col-0, (2) *Arabidopsis lyrata*, (3) *Brassica rapa*, (4) *Eutrema parvulum*, (5) *Aethionema arabicum*, (6) *Tarenaya hasslerania*, (7) *Carica papaya*, (8) *Citrus sinensis*, (9) *Vitis vinifera*, (10) *Solanum lycopersicum* and (11) *Solanum tuberosum*. Tandem duplicate gene copies are highlighted in red. Singleton genes are highlighted in dark blue. Conserved Cluster A-D refers to 4 distinct *A. thaliana NB-LRR* loci (A: AT3G14470; B: AT3G50950; C: AT4G33300; D: AT5G17860) including syntelogs if present in all other 10 genomes. Loose scaffolds and contigs not anchored to the genome assembly are shown shifted in radius but not in length scale. For genomes without assembly to the chromosome level, the 20 largest scaffolds are displayed. For genome assembly versions used in this analysis, see fig. 3. Please note that for due to fragmented assembly status of *Nicotiana benthamiana*, all scaffolds are below visible length threshold.

**Table 1.** Comparison of *NB-LRR* locus-containing Tandem Arrays* among 12 species.

|  | NBS-LRR genes | Tandem Duplicates<br>Number of | Fraction of Tandem<br>Duplicates | Tandem Arrays<br>Number of | Average Number of<br>Genes per Array | Number of Genes<br>in largest Array |
| --- | --- | --- | --- | --- | --- | --- |
| *B. rapa* | 167 | 92 | 55% | 31 | 2.9 | 8 |
| *E. parvulum* | 72 | 37 | 51% | 13 | 2.8 | 9 |
| *A. thaliana* Col-0 | 140 | 94 | 67% | 32 | 2.9 | 8 |
| *A. lyrata* | 166 | 71 | 43% | 23 | 3.1 | 9 |
| *Aet. arabicum* | 112 | 71 | 63% | 21 | 3.4 | 11 |
| *T. hasslerania* | 59 | 26 | 44% | 10 | 2.6 | 6 |
| *C. papaya* | 44 | 32 | 72% | 10 | 3.2 | 5 |
| *C. sinensis* | 455 | 136 | 30% | 61 | 2.2 | 5 |
| *V. vinifera* | 294 | 206 | 70% | 62 | 3.3 | 10 |
| *N. benthamiana* | 233 | 58 | 25% | 26 | 2.2 | 5 |
| *S. tuberosum* | 402 | 238 | 59% | 77 | 3.1 | 8 |
| *S. lycopersicum* | 219 | 125 | 57% | 40 | 3.1 | 7 |
| **Σ** | 2,363 | 1,186 | 50%* | 406 | 2.9 | 7.6 |

\* Difference of value compared to fig. 5 is due to presence on *N. benthamiana*

However, our data indicate that both aforementioned situations with eleven (*Aethionema*) and five (orange) genes-per-array are outliers beyond the average degree of *NB-LRR* gene tandem array extension. For example, the majority of all 1,191 tandem duplicates (60%) are organized in arrays with two genes only. Three gene members per array still occur in 18% of all cases, whereas four, five and more than five genes per array occur with a frequency below 10%, respectively (**Fig. 6**).

**Figure 6.**
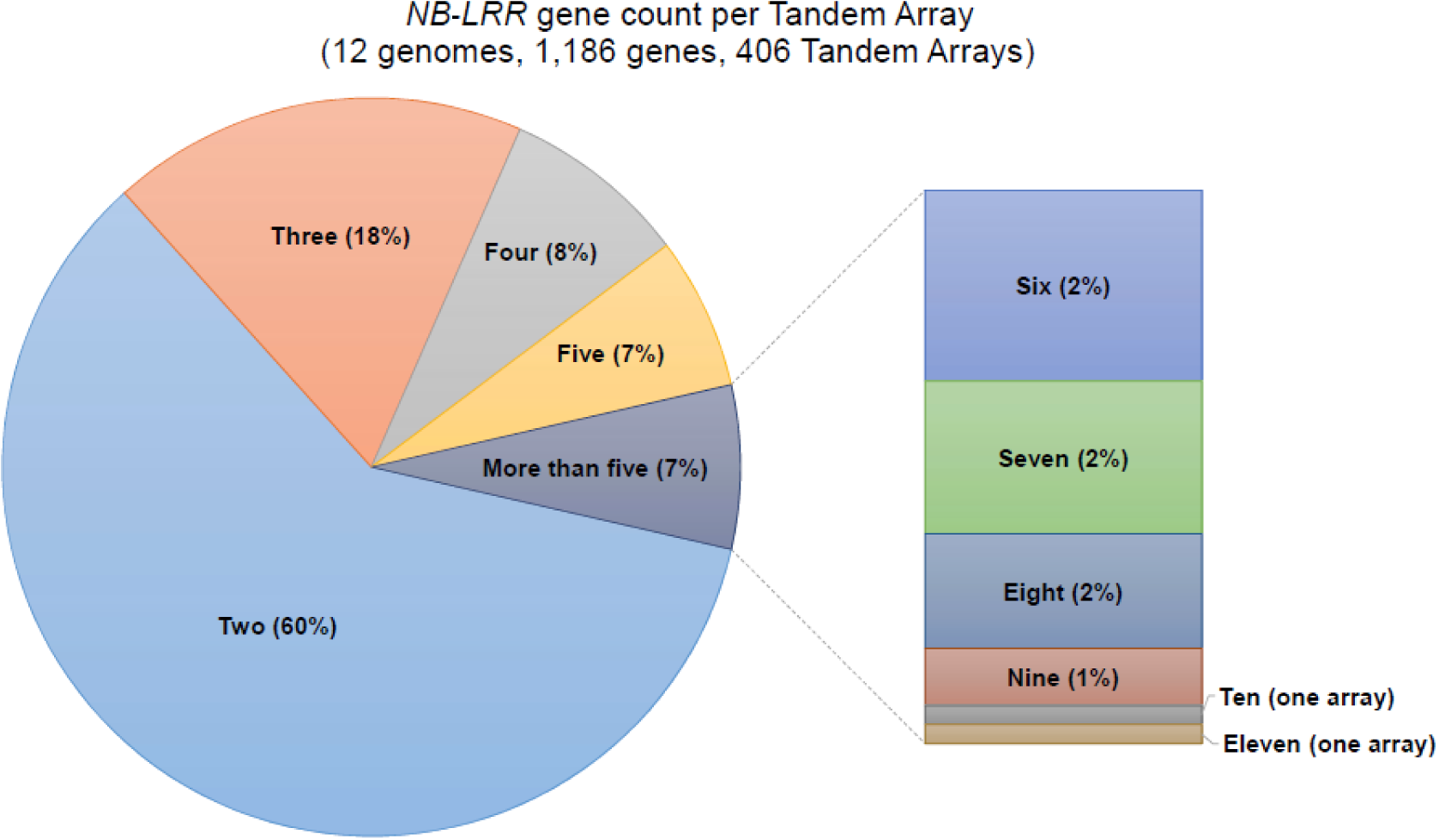
Gene count listing of full-length *NB-LRR* candidate gene-containing Tandem Arrays observed within all twelve analyzed genomes. 60% of all Tandem Arrays comprise two duplicate gene copies.

### Genome-wide determination of retained ohnolog duplicate fractions and utilization of R-proteins

Likewise, we determined the genome-wide fraction of retained duplicate groups due to ancient polyploidy events (ohnologs), including all *NB-LRR* loci. Screening of pairwise synteny blocks within the analyzed genome assemblies was accomplished using an integer programming approach implemented by the CoGe system for comparative genomics (see methods) (Tang et al. 2011). Due to technical restrictions, this was possible for seven genomes (i.e. minimum requirements in the N50 index, requiring a minimum of approximately 50 kb, see methods). The high degree of tandem duplicates among R-proteins in all species results in a low degree of retained ohnolog duplicates by definition, because ohnologs mainly comprise groups of two or three duplicates, whereas tandem arrays can have up to eleven members (**Fig. 6**). Notably, the *B. rapa* genome possesses the highest syntenic depth value among all analyzed genome assemblies with 12x in total (**Fig. 3**). Consistently, we found the highest fraction of retained ohnolog duplicates both genome-wide (53%) and among *NB-LRR* genes present in this crop with 42% in total. In contrast, the potato crop (*S. tuberosum*) contains the lowest fractions of retained ohnolog duplicates for both genome-wide average (10%) and the set of *NB-LRR* genes (5%), respectively. On average, ∽30% of all genes present in the seven analyzed genome assemblies comprise retained ohnolog duplicate groups. This fraction drops to 22% among all *NB-LRR* loci. This apparent under-representation of ohnologs among R-proteins highlights the high relative contribution of tandem duplication in R-protein cluster extension for the group of genome assemblies subjected to this analysis (Table 2).

**Table 2.**
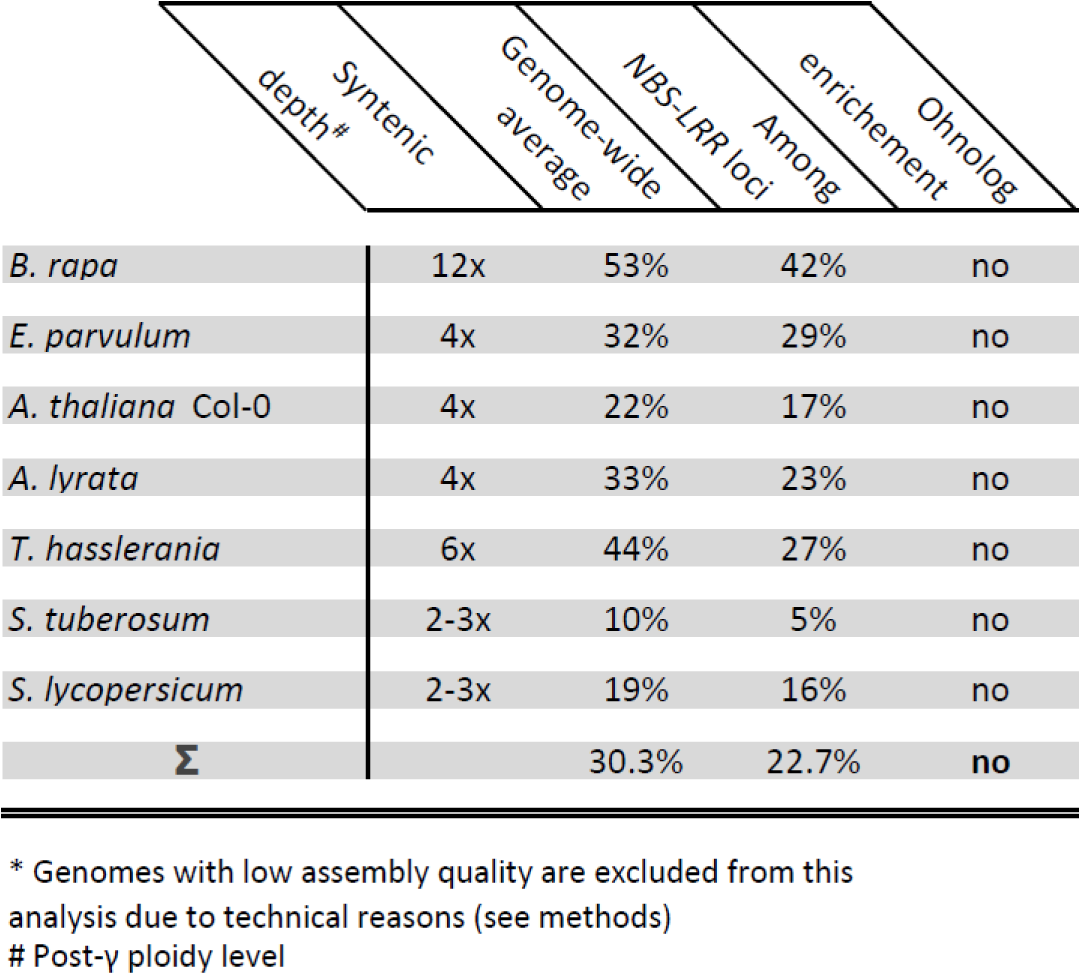
Species-wise comparison of retained ohnolog duplicates gene pairs among *NB-LRR* loci, shown for seven species*. Genomes with below-threshold mean and median scaffold size (N50 ∽50 kb) are excluded from this analysis due to technical reasons.

### Uncovering patterns of selection acting on the NB-ARC and LRR domain among all *NB-LRR* loci

We performed a genome-wide analysis of molecular evolution acting on all encoded *NB-LRR* proteins based on both the NB-ARC and LRR domains. In a first step, we grouped (a) members of tandem arrays, (b) retained ohnolog duplicates as well as (c) singleton genes (defined as non-tandem array genes without retained ohnolog duplicate). By analyzing non-synonymous substitutions per non-synonymous sites, compared to synonymous substitutions per synonymous site (Ka/Ks ratio or ω, dN/dS), patterns of strong positive selection were uncovered among all three groups. Strikingly, we also found differences in molecular evolution rates among all three groups. Members of tandem arrays evolved fastest with a ω mean of 1.59. In contrast, all analyzed retained ohnolog duplicates evolved with an intermediate rate (ω mean = 1.36). We report the slowest rate of molecular evolution for singleton *NB-LRR* genes with a ω mean of 1.22 (**Fig. 7**). Values for ω above one indicate positive or Darwinian selection, less than one implies purifying (or stabilizing) selection whereas ratios of one are indicative for neutral (i.e. absence of) selection (Hurst 2002).

**Figure 7.**
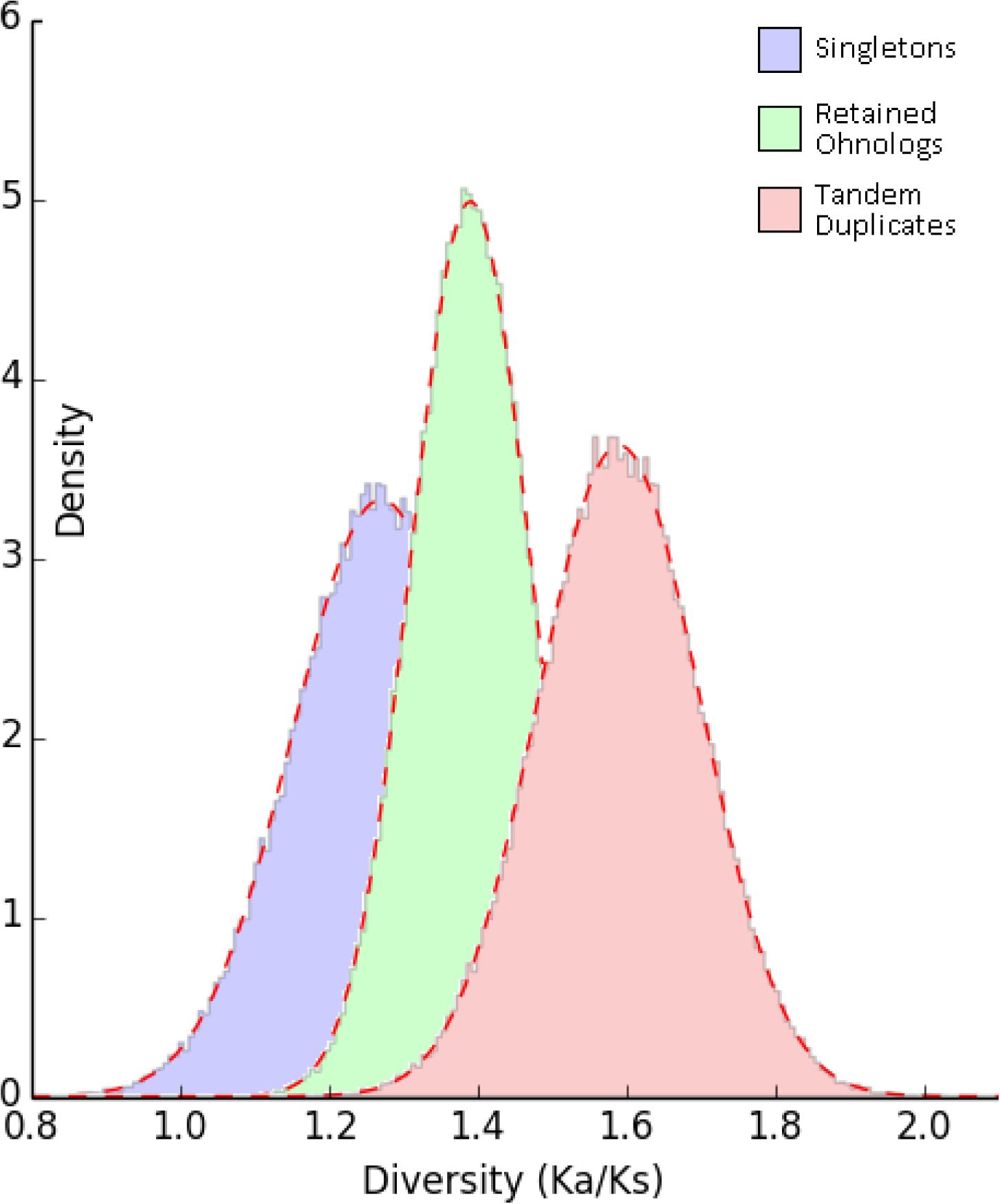
Selection in action between gene pairs of three major duplicates categories – singletons, tandem duplicates and WGD duplicates (ohnologs). Strong positive selection following gene and genome duplication of *NB-LRR* loci, as indicated by higher Ka/Ks values.

### Identification of *NB-LRR* functions conserved across all analyzed genome assemblies

Utilizing the wealth of *NB-LRR* functional and molecular data available in *Arabidopsis* as a reference, we composed a species-wide matrix of R-protein presence/absence based on sequence homology (i.e. filtered/non-filtered RBH) and synteny (**supplementary tables 1 and 2**). Among the extended set of 140 distinct *NB-LRR* functions encoded by the model plant (see above), we found four conserved clusters of “gatekeeper” genes sharing syntenic orthologs across all 12 analyzed genomes (**supplementary table 1**, **Fig. 5**). For two among those, functional data are available in *Arabidopsis*, whereas members of the other two gene clusters have not yet been characterized in any of the analyzed plant lineages.

The non-TIR non-CC *NB-LRR* (NL) class R-protein AT3G14460 is a “gatekeeper” because it forms one of four conserved clusters together with all of its aforementioned syntelogs (**supplementary table 1**, “Conserved Cluster A” in **Fig. 5**). Interestingly, there are yet no functional data available concerning this gene, neither in *Arabidopsis* nor in any of the other 11 analyzed genome/gene-space assemblies.

For example, this NL-class “gatekeeper” AT3G14460.1 (Meyers et al. 2003; Tan et al. 2007) forms syntenic RBH pairs with fgenesh2_kg.3__1571 (*A. lyrata*), Bra027333 (*B. rapa*), Tp3g12770 (*E. parvulum*), AA_scaffold578_71 (*Ae. arabicum*), Th16129 (*T. hasslerania*), supercontig_77.89 (*C. papaya*), GSVIVT01013307001 (*V. vinifera*), Solyc03g078300.1 (*S. lycopersicum*) as well as PGSC0003DMG400005046 (*S. tuberosum*), respectively. For *C. sinensis*, the RBH partner orange1.1g000782 m is harbored by a very small scaffold (∼12.6 kb) with 3 genes only, making the scoring of gene synteny impossible (**Fig. 8**). However, the locus orange1.1g000782 m in turn forms RBH pairs with the aforementioned genes supercontig_77.89 (*C. papaya*) as well as GSVIVT01013307001 (*V. vinifera*), thereby closing the gap in a phylogenetic framework (data not shown). Likewise, the *N. benthamiana* gene NB00009911g0001.1 forms RBH pairs with the aforementioned syntenic orthologs in tomato and grape-vine, overcoming the lack of synteny data for this early-stage draft genome assembly (data not shown). Notably, the underlying locus underwent tandem duplication after grape-vine lineage split, leading to presence of a tandem array in all Brassicales including orange, but an evident singleton gene in Solanaceae and *V. vinifera*, respectively (**Fig. 8**).

**Figure 8.**
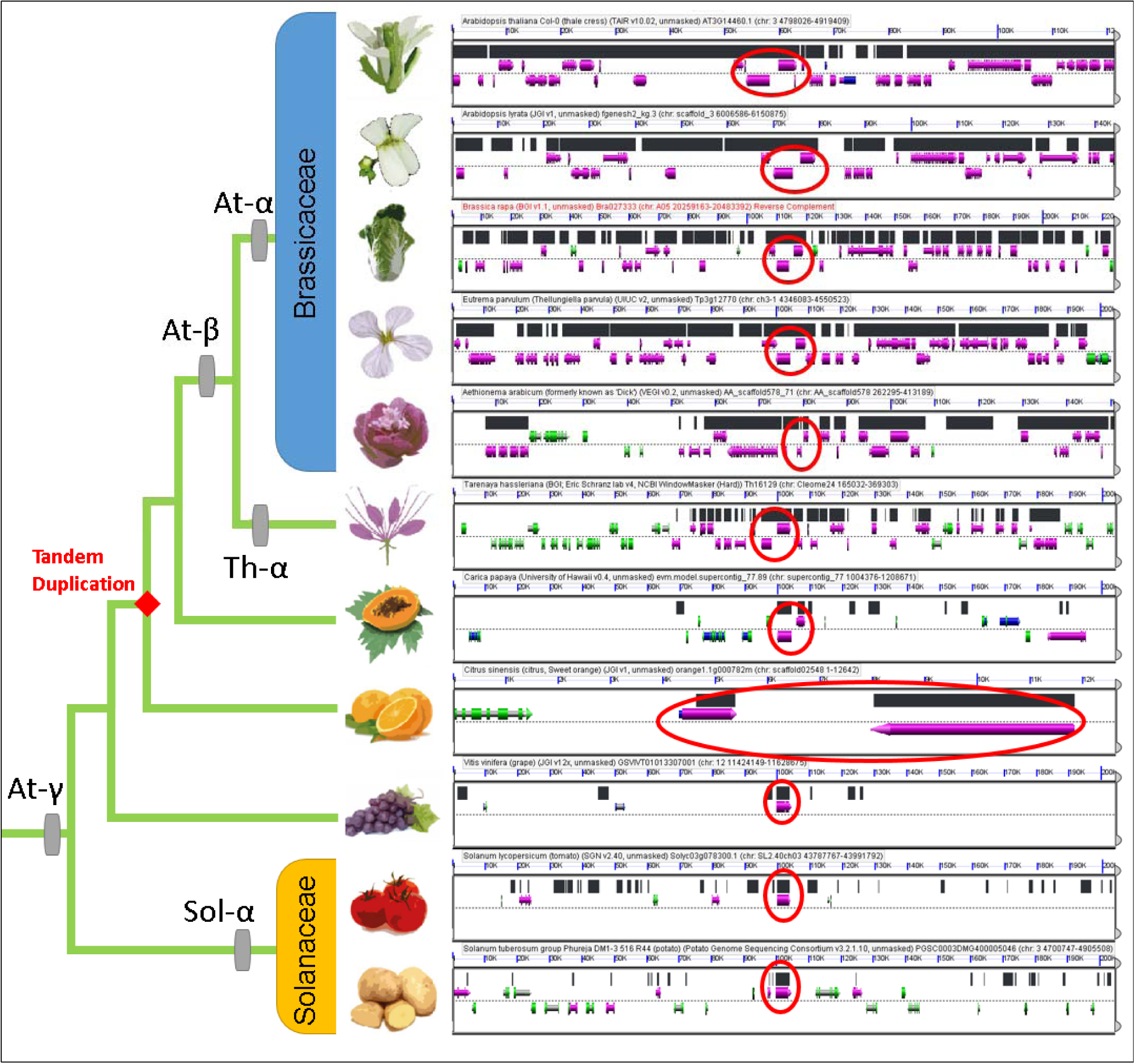
(B)LastZ 11-way multiple alignment of conserved cluster A from fig. 5. Shown left is phylogenetic relationship among all 11 species (*Nicotiana benthamiana* is excluded from this analysis due to technical reasons). Shown right is genomic context of the syntenic regions (marked in black). The regions in focus include one *NB-LRR* gene that expanded to a tandem array in the *Arabidopsis* lineage after split of Solanaceae. Diamond indicates tandem duplication event. Genes not overlapped by HSPs are shown in green. In case of *C. sinensis*, the orthologous genes is harbored by a very small scaffold (^∽^12.6 kb), therefore scaled differently from other panels in GEVO.

The *TIR-NB-LRR* (TNL)-class “gatekeeper” locus AT5G17680 is anchoring another group of syntenic orthologs shared by all lineages (**supplementary table 1**, “Conserved Cluster D” in **Figure 5**). Similarly, this locus lacks evidence on gene function in any of the analyzed plant lineages.

In contrast, conserved clusters B and C are anchored by *ZAR1* (HOPZ-ACTIVATED RESISTANCE 1 or AT3G50950) and *ADR1-L1* (ACTIVATED DISEASE RESISTANCE 1 or AT4G33300) in *Arabidopsis*, respectively (**supplementary table 1**, “Conserved Cluster B and C” in **Fig. 5**). *ZAR1* encodes a CC-NB-LRR (CNL) class R-protein of the FLARE gene group (Flagellin Rapidly Elicited, due to rapid up-regulation following exposure to the PAMP flg22) (Navarro et al. 2004). ZAR1 confers allele-specific recognition of the *Pseudomonas syringae* HopZ1a type III effector in *Arabidopsis* and acts independent of several gene products required by other R protein signaling pathways (Lewis et al. 2010). Likewise, *ADR1-L1* encodes a CC-*NB-LRR* (CNL) type R-protein with roles not only in defense response but also in drought tolerance; overexpression of *ADR1-L1* results in a dwarf phenotype and activation of defense-related gene expression in *Arabidopsis* (Navarro et al. 2004; Kato et al. 2011).

## DISCUSSION

The proliferation of higher throughput DNA sequencing and genome informatics approaches enables an accelerated production rate of draft genomes from a wide phylogenetic sampling of plant taxa, highlighting a need for robust methods and a comparative framework for gene and genomic comparisons. We therefore have developed a custom approach to identify functional groups of plant proteins applying a novel and highly complementary combination of available algorithms and datasets. We have applied this to R-proteins and annotated 2,363 loci of the *NB-LRR* type in total. This set contains genes that previously remained un-identified or under-explored for all species except tomato and potato. For Solanaceae, we stress that re-sequencing approaches based on complexity reduction such as target gene capture have been successfully applied for a similar purpose (referred to as Renseq) (Jupe et al. 2013; Uitdewilligen et al. 2013). However, it is not unreasonable to assume that the onset of next generation sequencing and genome informatics will continue with acceleration beyond Moore's law and hence lead to more and better algorithms for *de-novo* generation of gene annotations. Therefore, the added value of the computational pipeline shown in this study will rise with the same rate. For future references, we are working on customization of our approach to make it suitable for application to whole sequence scaffolds/contigs rather than sets of annotated genes/proteins. We intend to generate a computational pipeline for in-silico target gene capture based on scoring of combined hits outside the annotated gene-space within a size-window common to protein-coding genes, thereby overcoming the evident limitations of currently available algorithms for *de-novo* gene annotation (**Jupe F, personal communication**). The pipeline shown in this study represents the first step towards this goal.

We highlight three major findings in this study, including (a) higher contribution of tandem gene expansion in R-genes, (b) higher selection ratio in tandem duplicates compared to ohnologs and singletons and (c) stable orthologous clusters established for some R-genes that are likely to indicate a common functional constraint. R-genes typically show an unusually high turnover rate due to the strong selection to keep up with the biological arms-race with the plant pathogens (Mondragón-Palomino et al. 2002; Chen et al. 2010). Consequently, R-genes follow a different evolutionary trajectory than proteins inside large protein complexes or genes with regulatory roles (Freeling 2009). In this context, the added value of our study lies within the wide phylogenomics scope of the underlying approach. Although similar findings are available in *Arabidopsis*, monitoring dynamics underlying target gene evolution for approximately 100MA (corresponding to radiation time of the core eudicots) results in higher confidence of the results’ validity and thus decrease the error-rate of the underlying statements.

Since tandem duplicates represent the majority of the R-gene duplicates that typically have a higher turnover rate, and additionally most of the R-genes have experienced high birth-and-death rate due to the persistent arms-race with the evolution of pathogen target effectors, most R-genes should have a fairly limited cross-taxonomic coverage (Michelmore and Meyers 1998; Ratnaparkhe et al. 2011). However, some gene clusters do indeed get fixed much earlier in evolution, such as the four gene clusters that we have shown here to be conserved over 100 MA in most (if not all) core eudicot genomes. Could these gene clusters represent shared immunity responses to common pathogens? In addition, the genes in these clusters could also act as “helper *NB-LRR*s”, mediating signal transduction downstream of various different *NB-LRR* receptors for activation during effector-triggered immunity, thereby leveraging functional constraints as previously made evident for *ADR1* family in *A. thaliana* (Bonardi et al. 2011; Roberts et al. 2013). More studies need to be done in order to unravel gene function underlying the retention of these unusually ‘stable’ R-gene loci. This is stressed by the fact that (some degree of) functional evidence accumulated for two of our four *NB-LRR “*gatekeeper” functions in *Arabidopsis*; in one case a crosstalk to another key trait seems possible. In contrast, such data lacks for the other two “gatekeepers”, notably including one TNL class R-protein. We hypothesize significant potential for extension of gene functional data regarding all 4 “gatekeeper” loci, either by gene-for-gene resistance towards yet-undiscovered pathogen effectors or by facilitating effector-triggered signaling downstream of other *NB-LRR* genes similar to “helper *NB-LRR*s”. Notably, a combination of both scenarios is evident in *Arabidopsis* and hence not unreasonable to occur in other cases (see above).

We further highlight the need for “uniform” standards for comparative studies, such as the method we use in this study that is applicable but by no means limited to R-gene families. Our approach consolidates multiple tiers of evidence, including the basic protein similarity, domain compositions, and genomic context (synteny). The combined “meta-method” has better accuracy than most past computational pipelines using only one line of evidence. Uniform standards also ensure that our gene family member counts are directly comparable with one another, making in-depth studies of the expansion-contraction dynamics of gene families possible. Furthermore, our method allows efficient screening of genome assemblies for near-complete curation of multi-domain and multi-gene family clusters. In the case of *NB-LRR* type R-genes, the resulting raw data provide a detailed overview of nucleotide diversity among all target genes within and between twelve lineages covering the whole core-eudicot clade. Utilizing the wealth of functional data in *A. thaliana*, this leads to species-wise mapping (presence/absence) of every *NB-LRR* function characterized in the model plant. Notably, these data can be used by breeders to identify both target locus as well as small RNA sequence requirements for fast and efficient migration of resistance A to organism B using the emerging techniques of genome editing. For example, the particular *NB-LRR* gene conferring the desired resistance can be selected from our curated dataset followed by calculation of the smallest nucleotide distance (or closest related) target gene in the desired organism. The sequence of the small RNA(s) necessary for engineering of the aforementioned nucleases (see introduction) can be inferred accordingly in order to design a minimum set of experiments necessary and sufficient for gene-editing and thus generating an extended spectrum of resistance in any of the crop subjected to our analysis. Going beyond plant innate immunity, we provide data on a network of anchor genes present in all analyzed genome assemblies, thereby referencing orthologs and paralogs of every gene family present in the model plant *Arabidopsis*. We thereby excel future efforts to extract plant gene function, ultimately necessary for crop improvement and increased rates of global food production.

## METHODS

### Hardware resources and software prerequisites

All analysis were performed on a commercial Lenovo ultrabook, model Thinkpad X1 Carbon with 8GB RAM and Intel Core i7 3667U CPU (2 physical/4 virtual cores). The in-house developed perl and python scripts required perl (strawberry v5.18) and python (v2.7) libraries including bioperl (v1.6.910) and biopython (v1.63) modules, respectively. The iprscan_urllib.py-script for HMM-based domain annotation (see below) required SOAPy, NumPy and urllib python modules. For blast screens, we employed the stand-alone command line version of ncbi blast 2.2.27+ (ftp://ftp.ncbi.nlm.nih.gov/blast/executables/blast+/LATEST/, last accessed on #INSERT) (Altschul et al. 1990). For platform-independent coupling and parallelization of all employed scripts and programs, we wrote batch wrappers using the notepad++ editor (www.notepad-plus-plus.org, last accessed on #INSERT).

### Genome annotations

The Complete sets of representative genes and proteins for twelve genome annotations were downloaded using www.phytozome.net (last accessed on #INSERT) (Goodstein et al. 2012). We included *Arabidopsis thaliana* TAIR10 (Swarbreck et al. 2008), *Arabidopsis lyrata* v107 (Hu et al. 2011), *Eutrema parvulum* v2 (Yang et al. 2013), *Brassica rapa* v1.1 (Wang et al. 2011b), *Carica papaya* v0.5 (Ming et al. 2008), *Citrus sinensis* v1 (Xu et al. 2013), *Vitis vinifera* v2 (Jaillon et al. 2007), *Solanum tuberosum* v3.2.10 (Xu et al. 2011) and *Solanum lycopersicum* v2.40 (Potato Genome Consortium 2012). *Aethionema arabicum* v0.2 (Haudry et al. 2013) *Tarenaya hasslerania* v4 (Cheng et al. 2013) and *Nicotiana benthamiana* v0.42 (Bombarely et al. 2012) genome annotations were made available by the authors.

### Confirmation and extension of the *NB-LRR* multi-gene family in *Arabidopsis thaliana*

We obtained 138 *NB-LRR* genes from (Guo et al. 2011) and queried them against the TAIR10 *A. thaliana* genome annotation in a blast screen without e-value threshold (forward run). We extracted all target sequences and queried them back against the *A. thaliana* TAIR10 genome annotation with an applied target sequence maximum threshold of 2 (reverse run). After removal of self-hits, we scored loci as *NB-LRR* genes if they were part of the target sequence pool in the forward run, and aligned to a *NB-LRR* gene as defined by Guo et al. in the reverse run. We thereby created an extended set of *A. thaliana NB-LRR* loci.

### Determination of orthologous gene anchors

In a first step for large-scale *NB-LRR* gene identification, we determined reciprocal best blast hits (RBH) for both (a) protein and (b) coding DNA sequences between *A. thaliana* Col-0 and all other eleven genome annotations in a blast screen without e-value thresholds. Since *NB-LRR* loci can comprise up to 5 different domain types connected by partially conserved linkers, the RBH approach can result in false positives due to short but highly conserved highest-scoring sequence pairs (HSPs) in functionally non-relevant (i.e. structural) parts of the protein. Therefore, we developed a python script to discard RBH pairs with a query/target sequence length ratio below 0.5 and above 2.0, respectively. We determined (c) additional, length-filtered RBH pairs for these loci within the aforementioned length ratio scope to form a 3^rd^ line of evidence for orthologous gene detection.

### Syntelog / ohnolog determination

Calculation of pairwise syntenic blocks within and between genomes is based on integer programming (Tang et al. 2011) but implemented to an easy-to-use web interface termed CoGe platform for comparative genomics (www.genomevolution.org, last accessed on #INSERT) (Lyons et al. 2008). Within all genome assemblies, we determined genes sharing the same genomic context to counterparts in the *A. thaliana* Col-0 genome annotation (defined as syntelogs) using the DAGchainer (Haas et al. 2004) and Quota-Align (Tang et al. 2011) algorithms implemented to the “SynMap” function within CoGe. To mask noise generated by successive duplication(s) of ohnolog blocks, we applied Quota-Align ratios for coverage depth consistent with the syntenic depth calculated for each genome annotation. For merging of adjacent syntenic blocks, we applied a threshold of n = 350 gene spacers. For syntelog gene pairs, we calculated rates of synonymous substitutions (Ks-values) using CodeML of the PAML package (Yang 2007) implemented to SynMap and applied Ks-value thresholds for ancient WGD events as previously described (Bowers et al. 2003). For determination of within-species syntelogs (comprising ohnolog blocks due to autopolyploidy events), we proceeded similar with the difference that we queried the target genomes against themselves instead of against *Arabidopsis*, using the “SynMap” function within the CoGe platform for comparative genomics (parameters: relative gene order with a minimum cluster size of 5 genes and a maximum chaining distance of 20 genes). For the lineage-specific WGD events known from *B. rapa*, *T. hasslerania*, *S. tuberosum* and *S. lycopersicum*, we set maximum thresholds for Ks value averages of ohnolog blocks (1.5) to eliminate noise of recent duplication events. Due to minimum requirements on assembly quality that apply for usage of SynMap, it was not possible to determine the fraction of ohnolog duplicates for the current gene-space assemblies of *Aethionema, Carica, Citrus, Vitis* and *Nicotiana* with the available algorithms. Synteny of genes within and between lineages was visualized using the GEVO function implemented to the CoGe platform for comparative genomics (see above).

### Determination of anchor paralogs and generation of extended multi-gene family cluster pool

We defined the orthologous gene sets as sum of three groups of RBH pairs (first group: based on length-filtered protein pairs; second group: based on non-length-filtered protein pairs; third group: based on non-length-filtered CDS pairs; see above for length filter criteria). We merged the orthologous gene sets with the syntelog gene set to create a set of putative homologous loci anchoring all *A. thaliana* gene families in all other analyzed genome annotations (“anchor pool”). In a next step, we performed a blast search without e-value thresholds to query all homologous anchor genes against all twelve genomes to determine putative paralogs of the anchor gene set (forward run). We extracted all target sequences and queried them against the *A. thaliana* Col-0 TAIR10 genome annotation with a target sequence maximum threshold of 2 (reverse run). After removal of self-hits, we scored loci as *NB-LRR* if they align to any member of the extended *NB-LRR* locus cluster in *A. thaliana* (see above). We defined all members of this pool as anchor paralogs, if they are not present within the set of homologous anchor genes (see above), thereby creating a highly accurate super-cluster of *NB-LRR* genes across twelve genomes.

### Hidden Markov Modeling and prediction of protein domains

The above-mentioned extended multi-gene family cluster of *NB-LRR* genes is based on both sequence homology and genomic location of its members. However, we observed an erosion of synteny across lineages relative to their phylogenetic distance. Furthermore, DNA sequence homology decreases with phylogenetic distance due to wobble rules for the 3^rd^ codon position. Likewise, the protein sequence homology between distant multi-gene family members can decrease due to synonymous substitutions of amino acids belonging to the same chemical class (i.e. aliphatic, aromatic, indolic). Therefore, we applied a final filtering step to remove false-positives from the extended *NB-LRR* gene cluster pool across all genomes. Using the iprscan_urllib.py script provided by the European Molecular Biology Laboratory (EMBL, Heidelberg, Germany) (https://www.ebi.ac.uk/Tools/webservices/download_clients/python/urllib/iprscan_urllib2.py, last accessed on #INSERT), we queried every member of the extended *NB-LRR* cluster pool to 14 algorithms that apply Hidden Markov Models for (protein domain) signature recognition (BlastProDom, FPrintScan, HMMPIR, HMMPfam, HMMSmart, HMMTigr, ProfileScan, HAMAP, PatternScan, SuperFamily, SignalPHMM, TMHMM, HMMPanther and Gene3D) (Zdobnov and Apweiler 2001). We overcame the one-sequence-at-a-time limitation of the EMBL server by writing batch wrappers for 25x-fold parallelization. To form a second layer of control we additionally tested all target genes for an encoded LRR domain using the LRRfinder algorithm version 2.0 available at http://www.lrrfinder.com/ (last accessed on #INSERT) (Offord and Werling 2013). As a result, we mapped all protein domains present in the putative multi-gene family cluster onto their genes in less than a day, and discarded all false positive genes from the cluster (i.e. genes not coding for at least one cluster-common domain). Final referencing of proteins with both NB-ARC and LRR domains was performed using a multi-vlookup array function in MS excel 2013.

### Determination of tandem duplicate gene copies

To determine the fraction of tandem duplicate gene copies, we queried the complete protein annotation of every genome assembly against itself in a blast screen without any e-value threshold and filtered our final set of target sequences from above outside a window of n = 10 allowed gene spacers in both directions from the query sequence s previously described (Rizzon et al. 2006).

### Multiple protein alignments

To generate multiple alignments of protein sequences, the stand-alone 64-bit version of MAFFT v7 was employed (http://mafft.cbrc.jp/alignment/software/, last accessed in #INSERT) (Katoh et al. 2002). First, all *NB-LRR* proteins were aligned species-wise together with the HMM-generated consensus sequence of the NB domain (available at http://niblrrs.ucdavis.edu/At_RGenes/, last accessed on #INSERT) as well as the LRR domain (available at http://smart.embl.de/smart/do_annotation.pl?DOMAIN=SM00370, last accessed on #INSERT) using the command line mafft.bat --anysymbol --thread 4 --threadit 0 --reorder -- auto input > output. Mesquite v2.75 (http://mesquiteproject.org, last accessed on #INSERT) was used with multi-core preferences to trim MAFFT multiple alignments down to the NB and LRR domain blocks. Trimmed blocks were re-aligned using MAFFT with the command line mafft.bat --anysymbol --thread 4 -- threadit 0 --reorder --maxiterate 1000 --retree 1 –localpair input > output.

### Codon alignments and determination of substitution rates

Re-aligned NB and LRR domain blocks were transferred to codon alignments using the CDS sequence counterparts and the pal2nal.pl script v14 (Suyama et al. 2006) (http://www.bork.embl.de/pal2nal/distribution/pal2nal.v14.tar.gz, last accessed on #INSERT). Gaps were allowed but manually edited wherever necessary. We allowed unusual symbols and manually edited mismatches between CDS and protein sequences wherever necessary. Synonymous and non-synonymous substitution rates were determined using the “KaKs_Calculator” software (https://code.google.com/p/kaks-calculator/wiki/KaKs_Calculator, last accessed on #INSERT) (Zhang et al. 2006) including 10 substitution rate estimation methods (model averaging was applied). Divergence rates are generally determined between pairwise alignments of homologous sequences. For determination of average divergence rates among singletons (i.e. non-TD non-ohnolog loci), we aligned singleton *NB-LRR* loci with the best non-self blast hit among all singletons within one species. For determination of average divergence rates among retained ohnolog duplicates, we aligned all ohnolog *NB-LRR* loci with the best non-self blast hit among all ohnologs within one species. In case of ohnolog triplets, we only considered the highest-scoring sequence pair (HSP). For determination of average divergence rates among arrays of tandem duplicate *NB-LRR* genes, we aligned the first with the last member of every array, thereby covering the majority of all tandem arrays (see Results). In a control step, we determined average divergence rates for all pairwise combinations within the largest tandem array in every species and did not find significant deviations (data not shown).

### Generation and graphical editing of figures

Ideograms of plant chromosomes/scaffolds/contigs were generated using the circos package (http://circos.ca/, last accessed on #INSERT) (Krzywinski et al. 2009). Histograms and Venn-diagrams were generated using the matplotlib package (http://matplotlib.org/, last accessed on #INSERT). Other figures were generated with MS office and graphically edited using the GIMP package (http://www.gimp.org/, last accessed on #INSERT).

## ACKNOWLEDGEMENTS

I would like to thank Florian Jupe for his valuable input with proof-reading of the manuscript. Likewise, thanks go to Detlef Weigel and the whole BMAP team for their inspiration and discussions during the onset of this project. Finally, I am grateful to Mariam Neckzei for her help with graphical editing of the figures.

### DISCLOSURE DECLARATION

The authors hereby declare that there is no conflict of interest.

